# Suspension Electrospinning of Decellularized Extracellular Matrix

**DOI:** 10.1101/2024.01.26.577473

**Authors:** Sarah Jones, Sabrina VandenHeuval, Andres Luengo Martinez, Eric Burgeson, Shreya Raghavan, Simon Rogers, Elizabeth Cosgriff-Hernandez

**Author notes:** Corresponding Author: Dr. Elizabeth Cosgriff-Hernandez 107 W. Dean Keeton BME Building, Room 3.503D Austin, TX 78712.

## Abstract

Decellularized extracellular matrices (dECM) have strong regenerative potential as tissue engineering scaffolds; however, current clinical options for dECM are limited to freeze-drying its native form into sheets. Electrospinning is a versatile scaffold fabrication technique that allows control of macro- and microarchitecture. It remains challenging to electrospin dECM; which has led researchers to either blend it with synthetic materials or use enzymatic digestion to fully solubilize the dECM. Both strategies reduce the innate bioactivity of dECM and limit its regenerative potential. Herein, we developed a new suspension electrospinning method to fabricate a pure dECM scaffold that retains its innate bioactivity. Systematic investigation of suspension parameters was used to identify critical rheological properties required to instill “spinnability,” including homogenization, concentration, and particle size. Homogenization enhanced particle interaction to impart the requisite elastic behavior to withstand electrostatic drawing without breaking. A direct correlation between concentration and viscosity was observed that altered fiber morphology; whereas, particle size had minimal impact on suspension properties and fiber morphology. The versatility of this new method was demonstrated by electrospinning dECM with three common decellularization techniques (Abraham, Badylak, Luo) and tissue origins (intestinal submucosa, heart, skin). Bioactivity retention after electrospinning was confirmed using cell proliferation, angiogenesis, and macrophage assays. Collectively, these findings provide a framework for researchers to electrospin dECM for diverse tissue engineering applications.

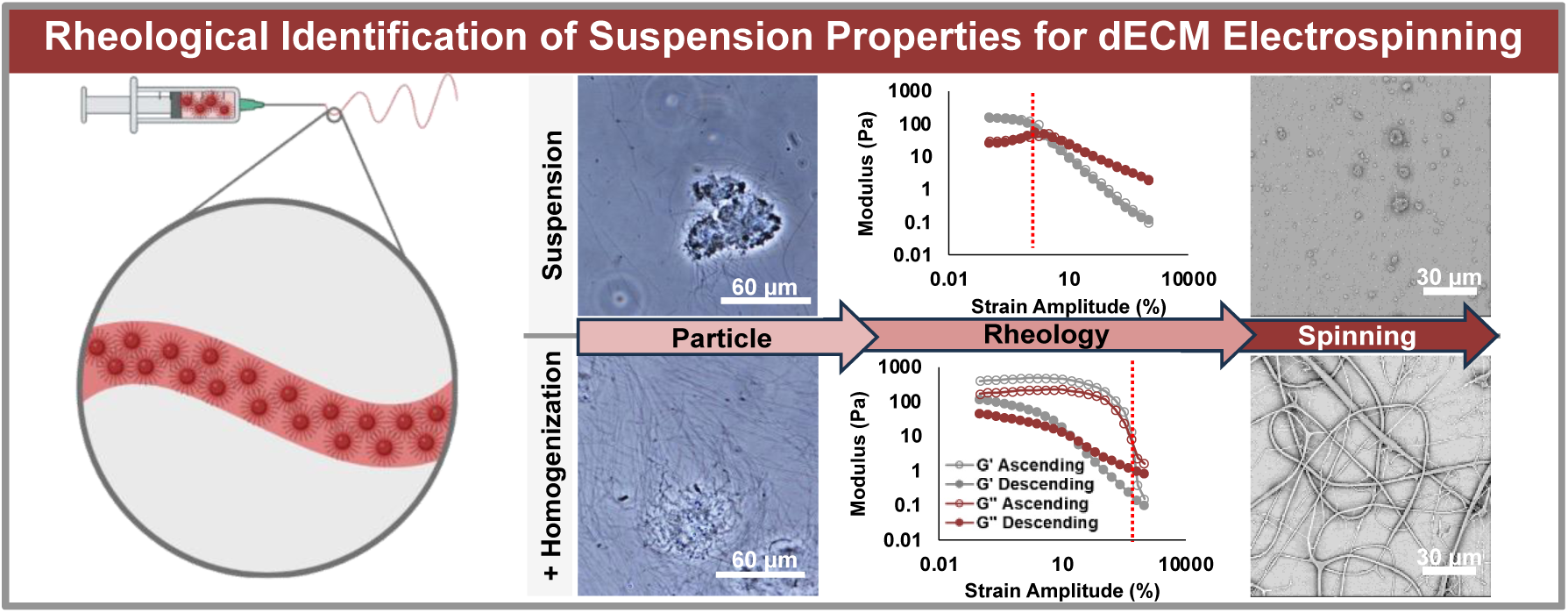

## 1. Introduction

Tissue engineering scaffolds composed of decellularized extracellular matrices (dECM) have shown a tremendous capacity to support regeneration in numerous applications.^1^ The dECM guides tissue remodeling by providing structural and biological cues with its complex composition of fibrous proteins, proteoglycans, growth factors, cytokines, and mRNA.^1–4^ Cells interact with and degrade the dECM, leading to the release of bioactive degradation products, such as chemoattractive low molecular weight proteins, angiogenic growth factors, and mRNA.^1, 4, 5^ Tissue-specific dECM has successfully promoted angiogenic, myogenic, neurogenic, and immunomodulatory behavior by supporting cell recruitment, proliferation, and tissue-specific differentiation.^6–12^ Small intestinal submucosa (SIS) is one of the most prominent dECM scaffolds on the market because it contains the high levels of growth factors and nutrients necessary to support the continual regeneration of the intestinal lining. SIS scaffolds have shown success in regenerating airway,^13^ abdominal wall,^14, 15^ diaphragm,^16^ intestine,^17^ bladder,^18^ rotator cuff,^19^ skin,^20–22^ and urethra^23^ tissue in clinical trials.^1^

Fabrication of dECM into specialized structures for a target application remains challenging. Bioactivity of proteins and growth factors can be easily deactivated with enzymatic digestion, heat exposure, or harsh solvents.^24^ Clinical practice is currently limited to native form sheets of dECM.^4, 7^ The macroscopic structure and microscopic morphology of dECM scaffolds are constrained to the structure of the harvested tissue, likely sheets or tubes. Current research strategies to expand the application of dECM scaffolds have been exploring hydrogels and combination devices like coatings and composites.^4^ Hydrogels provide a swollen matrix with potential for injection.^6, 25, 26^ However, gel fabrication requires enzymatic digestion to solubilize the ECM, which can denature and deactivate important components reducing the functional bioactivity of the dECM.^27, 28^ Similarly, dECM coatings and composites can have improved mechanical properties or structures, but they contain alternative materials that can alter the foreign body reaction and tissue remodeling response.^29–32^ A method is needed that can produce dECM scaffolds with retained bioactivity, ability to modulate geometry, and without additive materials that can produce negative biological responses.

Electrospinning is a fabrication platform that allows control of macro- and micro-architecture and can be scaled to fabricate meshes with clinically relevant dimensions. For example, electrospun materials can be fabricated with complex macrostructures such as valves, grafts, dressings, and wraps. Additionally, fiber microarchitecture can be controlled to vary anisotropy, pore size, and surface texture.^10, 33–38^ For instance, the fibers can be aligned to support regeneration of anisotropic tissues like skeletal muscle or randomly oriented to support regeneration of isotropic tissue like adipose tissue. The interconnected porosity and high surface area of electrospun scaffolds also provide a tunable platform for cell interaction and infiltration. Although electrospinning shows promise for dECM scaffold fabrication with high bioactivity retention and versatile structure, it remains common to use polymeric additives to facilitate fiber formation of dECM while electrospinning.^39–48^ Conventional electrospinning requires mobility of polymeric chains for alignment along the electrostatic field and chain entanglement to withstand the electrostatic forces.^49^ The polymeric additives provide increased chain entanglement; however, additives can negatively alter the biological response to the scaffold.^29, 30^ Other methods to enhance spinnability utilize enzymatic digestion to break down the ECM into individual chains, but this can reduce bioactivity through denaturation.^27, 28^ In contrast with traditional spinning solutions of digested and dissolved dECM, our recent work showed that a new suspension electrospinning method using decellularized skeletal muscle. The resulting scaffold retained myogenic bioactivity as indicated by cell attachment, proliferation, and differentiation.^10, 50^ However, electrospinning of dECM suspensions is in its nascency and thus critical rheological parameters to obtain a spinnable suspension have yet to be elucidated.^10, 50, 51^ A rigorous evaluation of suspension properties and its capacity for fiber formation is needed to expand the use of electrospun dECM scaffolds to a wide range of tissue engineering applications.

In this study, the critical properties of dECM suspensions that are required to facilitate electrospinning, or instill “spinnability,” are identified. We conducted a rigorous rheological evaluation of a wide range of dECM suspensions fabricated with varying homogenization levels, concentrations, and ground particle sizes with dECM derived from SIS. The versatility of the dECM suspension electrospinning method is showcased by electrospinning different ECM compositions, either prepared by different decellularization techniques or tissue origins. The level of bioactivity retention was then assessed via cell proliferation, macrophage polarization, and angiogenesis. This assessment of the critical parameters involved in electrospinning dECM aims to provide an accessible toolbox for research groups to create specialized dECM scaffolds with high bioactivity retention for numerous tissue regeneration applications.

## 2. Materials and Methods

### 2.1 Materials

All materials were purchased from Sigma Aldrich unless otherwise noted.

### 2.2 Small Intestinal Submucosa Isolation

The submucosal layer of a 6-9 mo porcine small intestine (Animal Technologies) was isolated as previously described.^52^ The small intestine was washed in PBS and cut into 10 cm sections. Each section was sliced longitudinally to form a flat sheet. The outer layer comprised of the serosa and muscularis externa was mechanically delaminated, and the inner mucosa layer was removed with repeated mechanical shearing. The remaining layer is the SIS. The SIS was stored in PBS at 4°C for no more than 24 h before beginning decellularization.

### 2.3 Tissue Decellularization

Each SIS decellularization technique explored has been named the Abraham,^53^ Badylak,^54, 55^ and Luo^6, 56^ technique based on the prior work. For the *Abraham technique*,^53^ 10 cm SIS sections were first submerged in 5 mL of 100mM ethylenediaminetetraacetic acid (EDTA) in 10 mM sodium hydroxide (NaOH) for 16 h. Second, the SIS was transferred to 5 mL of 1M hydrochloric acid (HCl) in 1M sodium chloride (NaCl) for 8 h. Third, the SIS was transferred to 5 mL of 1M NaCl in 1X PBS for 16 h. Fourth, the SIS was transferred to 5 mL of 1X PBS for 2 h. Lastly, the SIS was transferred to 5 mL of DI water for 2 h. All cell studies used SIS decellularized with the Abraham technique. For the *Badylak technique*,^54, 55^ 10 cm SIS sections were first submerged in 5 mL of 0.1% peracetic acid and 4% ethanol for 2 hours. Second, the SIS was transferred to 5 mL of PBS for 15 min. Third, the SIS was transferred to 5 mL of DI water for 15 min. Fourth, the DI water was exchanged for another 15 min. For the *Luo Technique,*^6, 56^ 10 cm SIS sections were each submerged in 5 mL of 1:1 methanol and chloroform for 12 h and subsequently washed with DI water. Second, the SIS was transferred to 5 mL of 0.05% Trypsin-EDTA for 12 h and subsequently washed with saline. Third, the SIS was transferred to 5 mL of 0.5% sodium dodecyl sulfate (SDS) in 0.9% NaCl for 4 h and subsequently washed in saline. Fourth, the SIS was transferred to 5 mL of 0.1% peracetic acid and 20% ethanol for 30 min and subsequently rinsed with saline. During each step of the three different decellularization techniques, the SIS was continuously agitated at 37°C. The decellularized SIS was stored at -80°C until further use. The native SIS was washed in 1X PBS without additional chemical treatment and stored at -80°C until further use. Decellularization following each technique was confirmed via DNA quantification. The DNA of the SIS was extracted following the manufacturer’s recommended protocol of the DNEasy Blood and Tissue Kit (Qiagen) and quantified with the Quant-iT™ PicoGreen™ dsDNA Assay Kit (ThermoFisher Scientific). There were 3 samples (n=3) in each group done in triplicate.

A 6-9 month porcine heart (Animal Technologies) was sectioned into 0.5 cm^3^ pieces and decellularized using the Badylak technique in 4 L of each medium described above with constant stirring. Skin tissue was isolated from Sprague Dawley rats. The hair was removed with Nair^TM^ Hair Remover. The skin was sectioned into 10cm^2^ pieces and decellularized using the Abraham technique in 5 mL/piece of each medium described above. The decellularized heart and skin was stored at -80°C until further use.

### 2.4 Preparation of dECM Suspensions

The frozen decellularized SIS sheets were submerged in DI water and thawed at 37°C. The SIS sheets were then homogenized with an OMNI Tissue Homogenizer (OMNI International). The homogenized SIS was flash-frozen in liquid nitrogen and lyophilized for 2 days. The dried SIS was broken into particles with an electric blade grinder (Kaffe) and separated into groups of < 250 µm, 250 – 500 µm, 500 – 2000 µm, and > 2000 µm with a sieve tower. SIS particles were suspended in chilled HFIP at 10 – 60 mg/mL concentrations and allowed to mix for over 12 h at 4°C. Immediately prior to electrospinning, the SIS suspensions were loaded into a capped syringe and homogenized for 20 sec. The frozen decellularized SIS sheets that were not homogenized were lyophilized immediately and then followed the same protocol as the dried SIS described above. The SIS suspensions that were not homogenized were transferred to a syringe and immediately electrospun. The decellularized skin suspension was prepared similarly with 500 – 2000 µm dried particle size, 70 mg/mL concentration, and both homogenization steps. The decellularized heart suspension was prepared similarly with 500 – 2000 µm dried particle size and 55 mg/mL concentration, and both homogenization steps.

### 2.5 Rheological Assessment

The dECM suspensions were analyzed on a Discovery Hybrid Rheometer, DHR-3 (TA instruments) with a 25 mm parallel plate geometry. Descending and ascending strain amplitude sweeps between 0.1 and 1000% strain were conducted at a constant angular frequency of 1.0 rad/s at 25°C to identify the amplitude dependence of the storage and loss modulus of the dECM suspensions. The crossover points of the storage and loss modulus curves were identified with linear interpolation. A flow ramp was conducted between shear rates of 0.1 to 1000 s^-1^ at 25°C to determine the rate-dependent viscosity of the dECM suspensions. There were 3 samples (n=3) in each group. A visual assessment of the particles in suspension contributing to the variations in rheological behavior was conducted. The SIS suspensions were diluted 10-fold to 4 mg/mL in HFIP. 20 µL of the diluted suspension was placed between 2 coverslips and imaged immediately before drying on a Nikon TS100 brightfield microscope. There were 3 samples (n=3) in each group and 5 images were taken of each sample.

### 2.6 Electrospinning

The dECM suspensions were pumped from a 20 G needle charged with 18 kV at a flow rate of 0.5 mL/h. A grounded copper plate covered in parchment paper was placed 20 cm from the needle tip. Electrospinning progressed until the mesh reached the target thickness. To prepare for imaging, the meshes were coated with 5 nm of gold (Sputter Coater 108, Cressington Scientific Instruments, Hertfordshire, UK). The meshes were subsequently visualized with scanning electron microscopy (SEM, Phenom Pro, NanoScience Instruments, Phoenix, AZ) at an accelerating voltage of 10 kV. There were 3 samples (n=3) in each group and 5 images were taken of each sample.

### 2.7 Cell Proliferation

The bioactivity retention of SIS after electrospinning was analyzed via cell proliferation. Electrospun SIS meshes and decellularized SIS sheets were cut into 8 mm discs and sterilized with UV irradiation for 15 min on each side. Human dermal fibroblasts (hDFs) were expanded in alpha minimum essential media (α-MEM) growth media, supplemented with 10% fetal bovine serum (Atlanta Biologicals, Flowery Branch, GA) and 1% of penicillin-streptomycin (10,000 U/ml, Thermo Fisher Scientific) at 37°C and 5% CO2. hDFs were seeded on the surface of the SIS samples at 25,000 cells/cm^2^. The cells were cultured for 1, 3, 5, and 7 days at 37°C and 5% CO2. The hDF metabolic activity was measured with the CellTiter 96 ® Aqueous MTS Kit (Promega Corp., Madison, WI, USA). At each timepoint, the samples were removed from the cell media and incubated with 17% (v/v) CellTiter 96 ® Aqueous One Solution Reagent in media at 37°C and 5% CO2 for 1 h. Subsequently, the absorbance of 100 µL of the media was analyzed on a plate reader (Infinite M Nano+, Tecan) at a wavelength of 490 nm. There were 3 samples (n=3) in each group done in duplicate.

### 2.8 Angiogenic Assays: In Vitro Tube Formation and Chorioallantoic Membrane (CAM) Assay

A tube formation assay was performed to evaluate the angiogenic capacity of the releasate from the SIS samples. Basal media (Endothelial Basal Medium 2 with 0.5% fetal bovine serum, Fisher Scientific) was conditioned with a decellularized SIS sheet, electrospun SIS mesh, and vascular endothelial growth factor (VEGF). Non-conditioned basal media was used as a negative control. Dried 5 mg SIS samples were sterilized with UV irradiation and soaked in 1 mL of basal media at 4C for 7 days with constant agitation. Recombinant human VEGF (R&D Systems) was added to the basal media at 20 ng/mL immediately prior to cell treatment. Human umbilical vein endothelial cells (HUVECs) were expanded in Endothelial Growth Medium 2 (Fisher Scientific) until confluent. The HUVECs were starved for 24 hours in basal media prior to the tube formation assay. Reduced Growth Factor Matrigel (Corning) was plated in a 96-well plate (50 µL per well) and incubated at 37C for 30 min. Aliquots of 400 µL were prepared for each media group containing 100 µL of the HUVEC suspension (40,000 cells/25 µL) and 300 µL of the conditioned media. Each media group was plated on the Matrigel at 40,000 cells per well and incubated at 37C for 4 hours. HUVECs were stained with Calcein-AM (Biotium) diluted to 1:2000 for 30 min and imaged with a fluorescent microscope (Nikon TS-100). The number of networks formed in each media group per image were manually counted. There were 3 samples (n=3) in each group and 5 images were taken of each sample.

A CAM assay was used to measure the angiogenic capacity of the SIS material as a whole. The samples tested include a decellularized SIS sheet, an electrospun SIS mesh, and a 200 µm Nylon mesh. Fertilized Japanese quail eggs (Southwest Gamebirds) were incubated horizontally at 37°C at 50% relative humidity for 3 days. The eggshells were sterilized with 70% ethanol and allowed to dry. The embryo was removed from the shell and placed in a 50 mm petri dish. Three 50 mm polystyrene dishes were placed in a 150 mm petri dish filled with 5 mL DI water to maintain a 100% humid environment and incubated at 37°C for 7 days. The samples were cut to 6 mm discs and placed 2 – 3 mm to the side of the central anterior vitelline vein in the outer third of the CAM as previously described.^57^ Images were taken daily for 4 days with a stereoscope to visualize angiogenesis. Vessel density was quantified as the change in the number of vessels crossing the outer perimeter of the sample from Day 0 to Day 4. There were 5 samples (n=5) in each group.

### 2.9 Macrophage Assay

Gene expression of macrophages cultured on SIS samples was analyzed to evaluate the retention of immunomodulatory bioactivity.^58^ THP1 monocytes (American Type Tissue Collection, Manassas, VA, USA) were expanded with non-adherent culture practices in RPMI media with 10% fetal bovine serum in an upright T-25 flask. THP1 monocytes were differentiated into M0 macrophages with 5 ng/mL phorbol 12-myristate 13-acetate (PMA) for 72 hours. A decellularized SIS sheet and electrospun SIS mesh cut to 5 mm discs were secured to a PDMS-coated well plate. Macrophages were seeded on the SIS samples at 1×10^5^ cm^2^ and cultured for 72 hours at 37C. Control macrophages were plated directly on the PDMS-coated well plate. RNA was extracted using an RNeasy Mini Kit (Qiagen) and the concentration was evaluated with a NanoDrop OneC (ThermoFisher Scientific). A High Capacity cDNA Reverse Transcription Kit (ThermoFisher Scientific) was used to perform reverse transcription. qPCR was used to detect the expression of NOS2, IL12, CD206, IL10, CHI3L1, VEGF, and TGFβ with CFX96 Real-Time System (Bio-rad) using the Applied Biosystems PowerSYBR Green PCR Mastermix (Thermofisher Scientific) for detection. The 2ΔΔCt method was used to calculate gene expression with GAPDH as the housekeeping control. The fold change in gene expression was calculated with respect to the control macrophages and the decellularized SIS sheet.

## 3. Results and Discussion

Based on our prior work electrospinning suspensions of decellularized skeletal muscle, the SIS suspensions were originally prepared by the Abraham decellularization technique, lyophilization, grinding into powder, and suspension in HFIP.^10, 50^ Regardless of concentration or particle size, the dECM collection was in the shape of spherical beads, indicating fiber break up. The viscosity of polymeric electrospinning solutions has been widely used to predict spinnability and fiber morphology.^59, 60^ Although the viscosity of the suspensions was between 1 and 4 Pa·s, which has previously been shown to be within the electrospinning range of 0.1 to 50 Pa·s, this did not yield fibers when spun.^49, 59, 61^ Increasing the concentration of the suspension to surpass a viscosity of 4 Pa·s and circumvent beading resulted in swollen particles without a liquid phase. The dECM suspensions exhibited less elastic rheological behavior than polymeric electrospinning solutions. The chain entanglements in polymer solutions instill elastic properties that resist fiber breakup during electrostatic drawing of the electrospinning process. We hypothesized that the lack of interaction between distinct particles did not provide sufficient elastic behavior within the suspension. In order to increase entanglement and elasticity, the dECM suspensions were homogenized. A stepwise approach was taken by homogenizing the native form dECM sheet prior to drying and/or homogenizing the dECM suspension immediately prior to electrospinning. Each suspension was prepared with a dried particle size of 500 – 2000 µm and concentration of 40 mg/mL. An extensive assessment was conducted to identify the structural changes within the suspension and the rheological properties that imparted “spinnability.”

### 3.1 Effect of Homogenization on Particle Interaction and Fiber Formation

The dECM suspensions were diluted 10-fold and imaged with brightfield microscopy to assess the structural change in response to homogenization **(Figure 2A)**. Overall, the particles in suspension are significantly smaller than the dry ground particles used to prepare the suspension. The prepared dry particles are 500 – 2000 µm, and the hydrated suspended particles are approximately 50 µm even though SIS has a high swelling ratio of around 2000%. This indicates that the dried particles are breaking down during the mixing of the suspension but are not dissolving. Additionally, the suspensions contain fibrillar components on the order of microns similar to the scale of collagen bundles.^62^ This indicates protein microstructure above the tertiary organization level is being retained. The particles within the suspensions fabricated with homogenized SIS sheets are less dense and exhibit large fibrillar networks protruding from their core. The surrounding fibrillar networks are not distinct between particles, rather they interact with fibrils protruding from other particles or fibrils that are freely suspended. The particles within the suspensions fabricated with non-homogenized SIS sheets do not exhibit large fibrillar networks, but rather short fibrils protruding from their core, if any. The homogenization of the SIS sheets has a greater impact on the density of the particles and the size of fibrils extending from their core, whereas the homogenization of the suspensions has a greater impact on the concentration of free fibrils in suspension not anchored by a particle. We hypothesize that the homogenization of the SIS sheet has a greater effect on the elasticity of the suspension because the initial density of the dried particles suspended in HFIP is reduced. This allows loosening of the particles, elevated formation of protruding fibrils, and increased particle-particle interaction. To evaluate this hypothesis of particle interaction facilitating elastic behavior, a rheological assessment was conducted.

**Figure 1:**
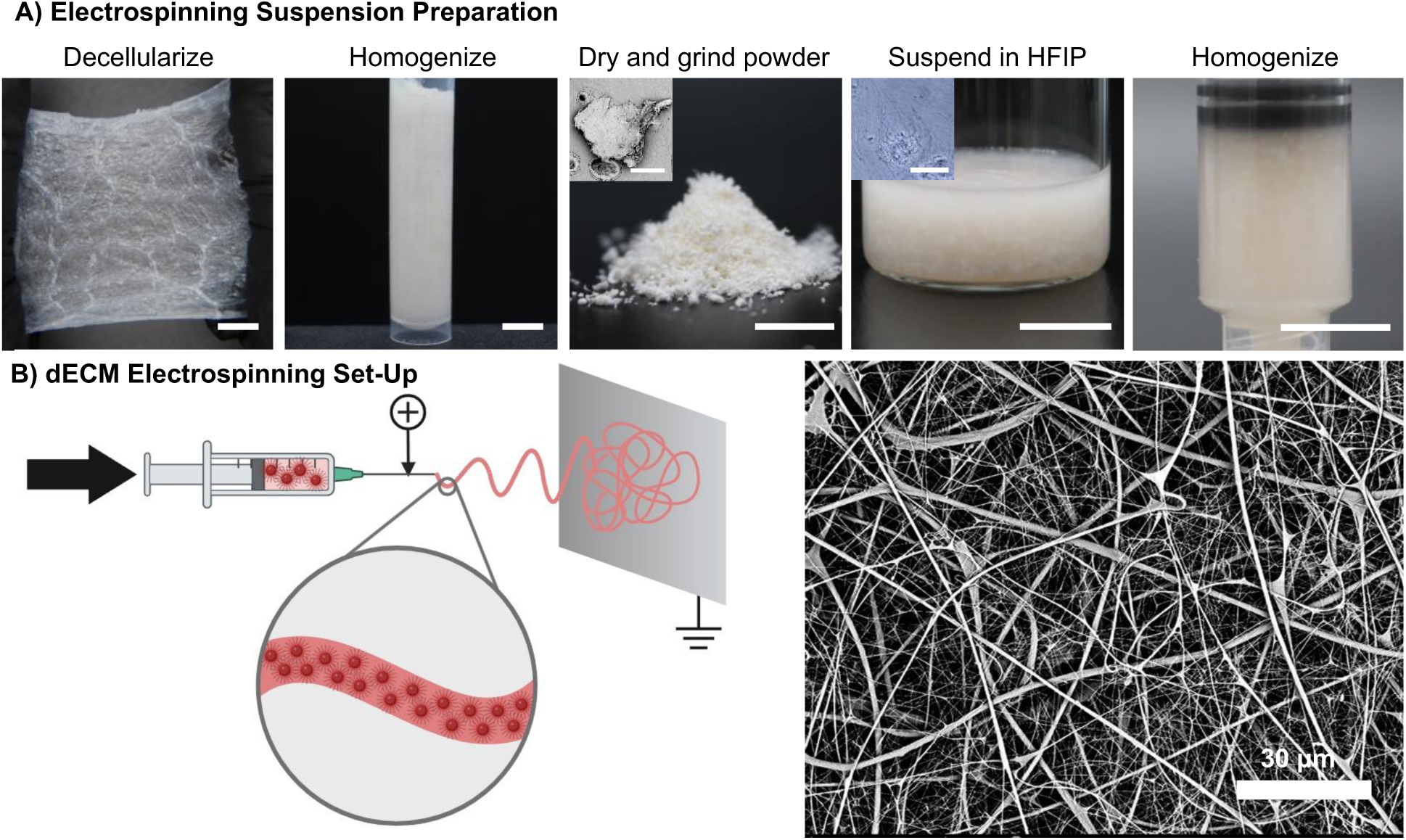
Schematic of electrospinning method for dECM. A) The preparation of a dECM electrospinning suspension is a six-step process involving decellularization, homogenization, milling, suspension in HFIP, and a second round of homogenization. Macro images: scale bar = 1 cm. SEM image of dried SIS particle: scale bar = 300 µm. Microscopic image of hydrated SIS particle in suspension: scale bar = 60 µm. B) The dECM suspension is electrospun by applying an electric field between the flowing suspension and a collector. Created in BioRender. SEM image of electrospun dECM fibers. Scale bar = 30 μm.

**Figure 2:**
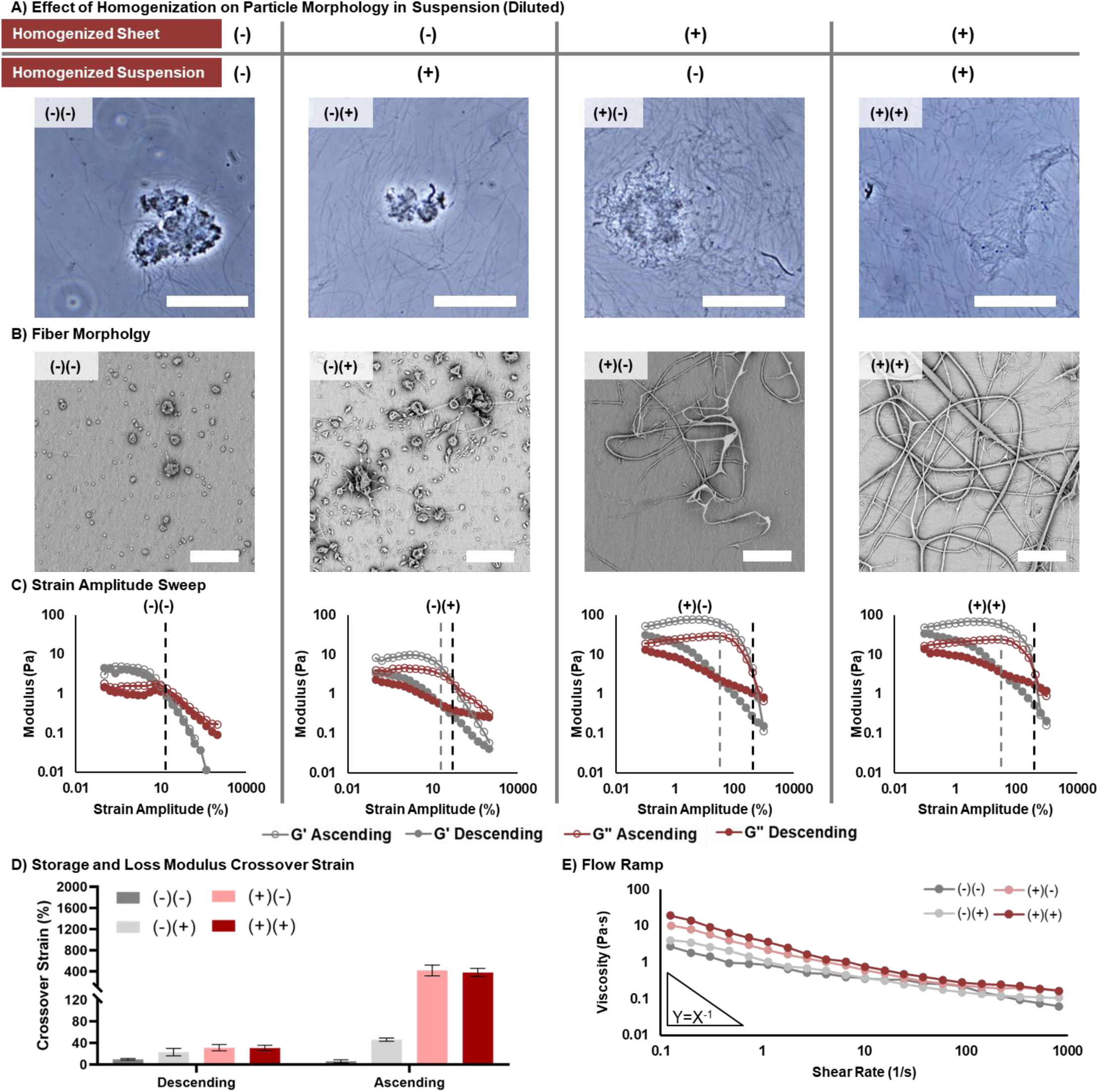
The effect of homogenization on suspension morphological and rheological properties. A) Brightfield image of dECM particles in an electrospinning suspension. Scale bar = 60 µm. B) SEM images of collection morphology. Scale bar = 20 µm. C) Strain amplitude sweep of dECM suspensions in which solid points were collected with a descending amplitude sweep and hollow points were collected in an ascending manner. The grey dotted line indicates the crossover from the descending amplitude test and the black dotted line indicates the crossover with ascending applied strain. D) The strain amplitude at which the storage and loss moduli cross collected with a descending strain sweep. E) The rate dependence of the viscosity of the dECM suspensions.

A strain amplitude sweep was conducted to assess the relationship between the storage and loss modulus of the suspensions across a range of strain amplitudes **(Figure 2C-D)**. In all cases we observe that the suspensions are viscoelastic solids at small amplitudes, as noted by the dominance of the storage modulus over the loss modulus, and that the suspensions are viscoelastic liquids at large amplitudes, where the loss modulus is larger than the storage modulus. While the transition from primarily elastic behavior to primarily viscous behavior has been shown to be complex and transient, a good measure of the yield point is the crossover between the storage and loss modulus. Above the crossover strain the suspension is flowing unrecoverably, while below the crossover strain the suspension acquires more strain recoverably than unrecoverably. Homogenization of the suspensions had a direct effect on the crossover point. The suspensions that underwent increasing homogenization had a larger crossover strain amplitude, and the homogenization of the SIS sheet had a greater impact than the homogenization of the suspension. The amplitude sweep was conducted in both ascending and descending fashion to determine if the rheological behavior of the suspensions is time-dependent, or thixotropic. The suspensions that underwent increasing homogenization exhibited a larger hysteresis loop indicating increased thixotropic behavior. Similar responses have previously been observed in hydrogels.^63^ The homogenization of the SIS sheet had a greater impact on thixotropy than the homogenization of the suspension. Lastly, a flow ramp was conducted to evaluate the rate-dependence of the viscosity of the suspensions **(Figure 2E)**. Although viscosity alone could not predict spinnability of suspensions, it is an important parameter to assess the full rheological profile of the suspensions. At shear rates greater than 10 s^-1^, all the suspensions had similar viscosities. Conversely, at shear rates less than 10 s^-1^, the viscosity increased with increasing homogenization, and there was a similar impact of suspension homogenization and SIS sheet homogenization. It was concluded that homogenization increased the elastic and thixotropic behavior of the dECM suspensions. This is indicated by increased crossover strain amplitudes and time-dependent responses. We hypothesize that the loose fibrillar components of the particles in suspension interact and produce drag between particles which is represented by increased elasticity of the suspension. When homogenized suspensions were tested with an ascending strain amplitude, the interaction of the particles overcomes the applied deformation until a sudden breaking point when the particles separate and the suspension flows. When decreasing the strain amplitude, the particles are initially separated by the large strains and are allowed to slowly establish interactions more gradually. Conversely, the particles within non-homogenized samples can slide past each other similarly regardless of the applied strain history because they lack entanglements. The spinnability of the established suspensions was assessed to identify the necessary rheological properties to fabricate a fibrous mesh.

The spinnability of the dECM suspensions was correlated to their rheological properties. With increased homogenization of the suspensions, a transition from beads to fibers was seen **(Figure 2B)**. The homogenization of the SIS sheet had a greater impact than the homogenization of the suspension. The suspensions fabricated from a non-homogenized SIS sheet created a bead-like morphology, and homogenizing the suspension led to minimal formation of strings branching from the beads. The suspensions fabricated from a homogenized SIS sheet created a fibrous morphology, and homogenizing the suspension led to more continuity within the fibers. Thus, the crossover strain and level of fibrils protruding from the particles in suspension have a greater impact on spinnability than the free fibrils in suspension. This suggests that the primary mechanism of fiber formation during spinning is the interaction between particles rather than the free fibril network. The fibril network is likely too dilute to withstand the electrostatic forces involved in electrospinning. However, the fibril network is expected to support the continuity between occasional disparate particles as showcased by broken fibers electrospun with the non-homogenized suspension and continuous fibers electrospun with the homogenized suspension. As expected, the strain amplitude at which the dynamic moduli cross when increasing the amplitude is more predictive of fiber formation than the descending crossover point because it is more representative of the increasing strain applied during electrospinning. The suspension begins unstrained in the syringe and is passed through a needle, during which significant amounts of strain are acquired. Upon leaving the needle, the suspension is drawn and whipped through the air with maximal strain before resting on the collector in a dried fibrous state. The suspensions with an ascending crossover strain amplitudes ∼ 400% resulted in fiber formation, whereas suspensions with an ascending crossover strain amplitude < 100% resulted in bead formation. Although the ascending crossover strain amplitude does not fully represent the complex rheological properties that are involved in electrospinning, it is a functional predictor of spinnability that assists in the high throughput preparation of dECM suspensions. A discrete crossover strain amplitude threshold was not identified with this stepwise homogenization approach, but a range of success was evaluated by assessing the impact of alternative suspension preparation parameters that alter rheological properties and fiber morphology.

### 3.2 Effect of Concentration on Rheological Properties and Fiber Formation

The impact of solution concentration on electrospun fiber morphology has been extensively reported.^49, 59, 64–68^ Surface tension and the applied electric field overcomes the chain entanglements in low concentration polymeric solutions causing them to break into beads. At higher concentrations, the chain entanglements can withstand the electric field and draw into a fiber. Suspensions do not follow the same chain entanglement paradigm; rather, they withstand electrostatic forces via particle interaction. Thus, the impact of concentration on suspension electrospun fiber morphology was evaluated **(Figure 3)**. SIS suspensions were prepared with concentrations from 10 – 60 mg/mL and homogenization of both the SIS sheet and SIS suspension (dual homogenized). Each suspension was prepared with the Abraham decellularization technique and a dried particle size of 500 – 2000 µm. The ascending crossover strain amplitude of the 10 – 50 mg/mL suspensions were all similar and greater than 400% **(Figure 3B-C)**. This indicates the suspensions are viscoelastic solids and deform elastically at high strains. Interestingly, the 60 mg/mL suspension exhibited a significantly reduced crossover strain amplitude compared to the 10 – 50 mg/mL suspensions. A flow ramp indicated that the viscosity generally increases with increasing suspension concentration, especially at high shear rates **(Figure 3D)**. Increasing the concentration of the suspension increases the density of particles and thus particle interaction. Under shear, increased particle interaction causes a suspension to have elevated resistance to deformation. However, the applied force at which these interactions break remains the same, as seen with the similar crossover strain amplitude required to make the suspension flow. Once electrospun, the 10 – 50 mg/mL suspensions all created fibers; however, fibers spun with 10 – 20 mg/mL suspensions contained beads while fibers spun with 30 – 50 mg/mL suspensions did not **(Figure 3A)**. Thus, the crossover strain amplitude remains a good predictor of spinnability and fiber formation, but viscosity is also needed to fine tune fiber morphology. The limited particle interaction in low concentration suspensions is overcome by the electrostatic forces in electrospinning and these suspensions begin to break into beads, as often occurs in dilute polymeric solutions due to limited chain entanglement. Additionally, electrospinning the 10 mg/mL suspension creates a largely fused mesh because the elevated solvent content is unable to sufficiently evaporate before collecting. There was no collection when the 60 mg/mL suspension was electrospun, because the viscosity of the 60 mg/mL suspension was too high for the electrostatic forces to deform. Voltage was applied up to 30 kV and no taylor cone was seen. It is expected that a higher electric field would be able to draw the 60 mg/mL suspension into a fiber. Altogether, the observed trends regarding concentration and electrospun fiber morphology of dECM suspensions remain consistent with electrospinning polymeric solution literature.^49, 69^ Thus, the impact of the concentration of particle interaction in suspensions is analogous to chain entanglement in solutions.

**Figure 3:**
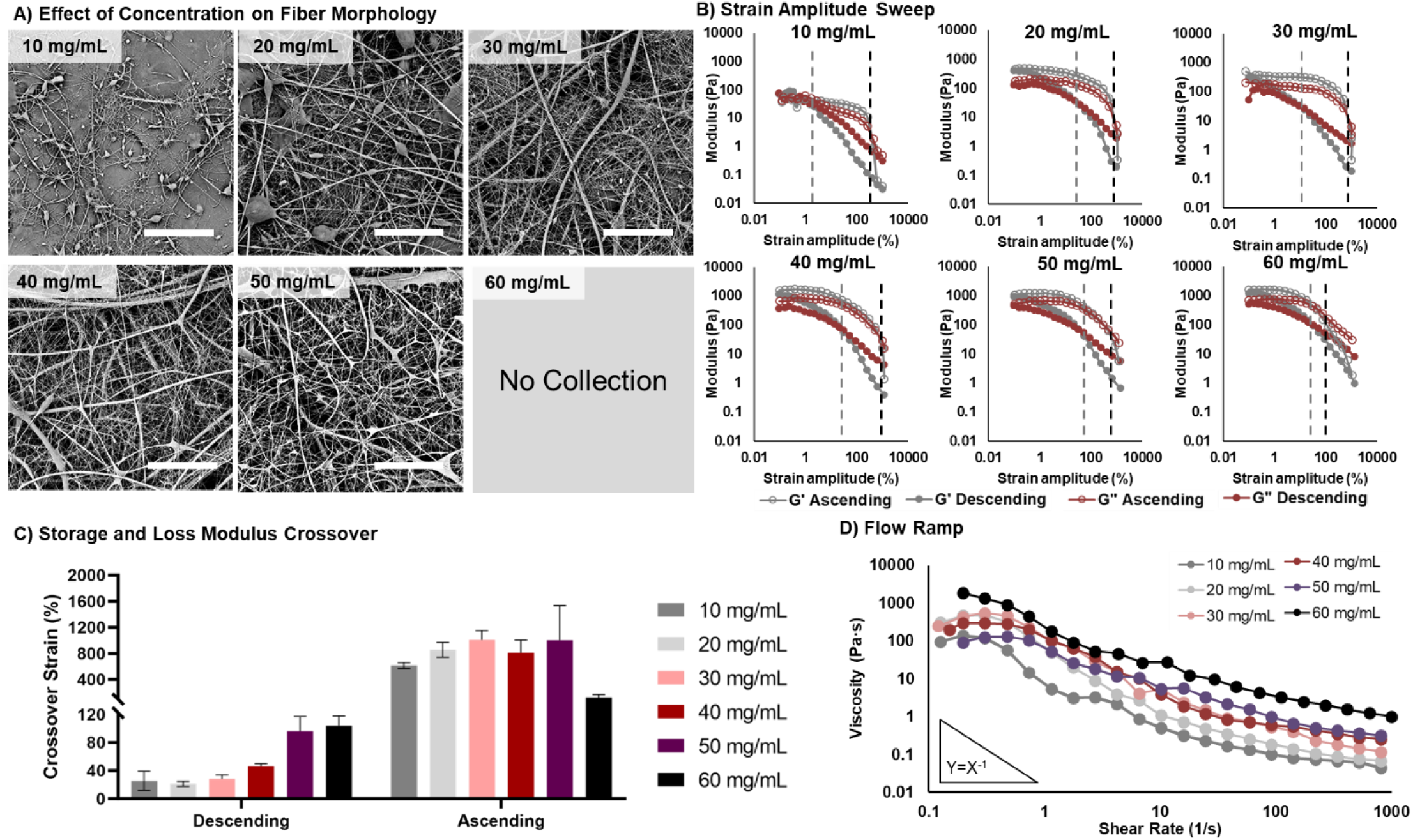
The effect of SIS concentration on suspension rheological properties and fiber morphology. A) SEM images of fiber morphology. Scale bar = 20 µm. B) Strain amplitude sweep of dECM suspensions in which solid points were collected with a descending strain sweep and hollow points were collected with an ascending strain sweep. The grey dotted line indicates the crossover with descending applied strain and the black dotted line indicates the crossover with ascending applied strain. C) The crossover strain of the storage and loss modulus curves collected with descending and ascending strain sweeps. D) The viscosity with respect to shear rate of the dECM suspensions.

### 3.3 Effect of Particle Size on Rheological Properties and Fiber Formation

A parameter controlled in dECM suspension preparation that is not applicable to polymeric solutions is dried particle size. Polymeric pellets completely dissolve in the chosen solvent; therefore, the initial pellet size does not impact solution properties. However, in dECM suspensions, the input dried particles do not dissolve in the suspension; rather, they remain as particles. The suspended particles (∼ 50 µm) are an order of magnitude smaller than the input dry particles (500 – 2000 µm) as seen in **Figure 2**. This suggests that the dried particles are breaking down into smaller constituents in suspension. The suspended particle size is expected to impact particle interaction and suspension rheological properties because size will affect the particle’s surface area. Thus, the impact of dried particle size on suspension particle morphology and electrospun fiber morphology was evaluated **(Figure 4)**. In this study, the dual homogenized SIS suspension was prepared with four size ranges of dried SIS particles including < 250 µm, 250 – 500 µm, 500 – 2000 µm, and > 2000 µm. Each suspension was prepared with the Abraham decellularization technique and a concentration of 40 mg/mL. Interestingly, the dried particle size from < 250 µm to > 2000 µm did not significantly impact suspension particle morphology **(Figure 4A)**, electrospun fiber morphology **(Figure 4B)**, or suspension dynamic moduli **(Figure 4C)**. At low shear rates, there was a minimal decrease in suspension viscosity with increasing dried particle size, but at high shear rates, there was not a significant impact of particle size. Each suspension is composed of hydrated particles on the scale of tens of microns with protruding fibrils and free fibrils similar to the dual homogenized suspension deemed “spinnable” in **Figure 2**. These results suggest that the dried SIS particles input into the HFIP break down to a consistent equilibrium state during mixing, regardless of the initial size. In order to further test this hypothesis that suspension particle size impacts rheological properties, dried particles less than 50 µm would need to be prepared. However, for this functional assessment on the controls needed for preparing spinnable dECM suspensions, it has been determined that the size of the ground particles does not need to be precisely controlled between 250 and 2000 µm. Thus, expensive tissue grinding methods such as cryo-milling and bead-milling are not required for dECM suspension electrospinning.

**Figure 4:**
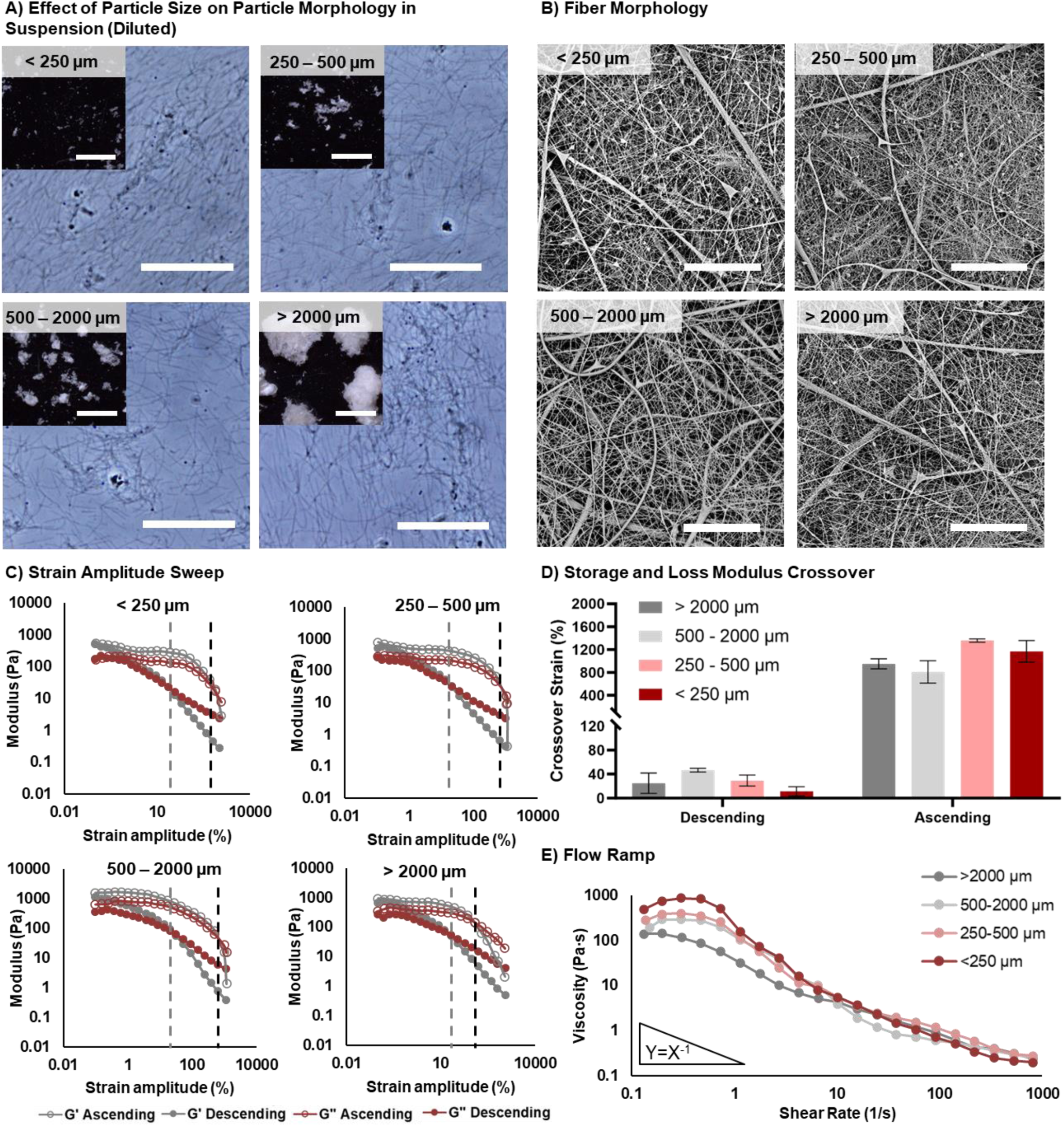
The effect of ground particle size on suspension rheological properties. A) Brightfield image of dECM particles in an electrospinning suspension. Scale bar = 60 µm. Inset stereoscope image of dried dECM particles. Scale bar = 3 mm. B) SEM images of fiber morphology. Scale bar = 20 µm. C) Strain amplitude sweep of dECM suspensions in which solid points were collected with a descending strain sweep and hollow points were collected with an ascending strain sweep. The grey dotted line indicates the crossover with descending applied strain and the black dotted line indicates the crossover with ascending applied strain. D) The crossover strain of the storage and loss modulus curves collected with descending and ascending strain sweeps. E) The viscosity with respect to shear rate of the dECM suspensions.

Altogether, a functional assessment has been conducted on the critical parameters needed to electrospin a dECM mesh with a suspension-based approach. Homogenization is critical to prepare a suspension that deforms elastically under elevated strain amplitudes and can withstand the electrostatic forces in electrospinning **(Figure 2)**. Instead of chain entanglement, dECM suspensions rely on particle interaction instilled by loose fibrillar networks protruding from individual particles. Homogenization reduces the density of the particles, increases the fibrillar component within the suspension, and thus increases the particle interaction and suspension elasticity. Once a spinnable dual homogenized suspension was prepared, the impact of concentration and particle size on fiber morphology was evaluated **(Figures 3-4)**. The dECM concentration in the suspension had similar impact as polymer concentration in solution, where lower concentration led to beading and higher concentrations overcome the capabilities of the applied field. Thus, particle interaction is analogous to chain entanglement with regards to spinnability of suspensions and solutions. Alternatively, suspension particle size does not have an analog in solutions because the polymer pellets completely dissolve. The dried particle size between 250 – 2000 µm did not have a significant impact on suspension properties or electrospun fiber morphology. The dried particles break down to a consistent size and morphology which suggests an equilibrium state of the hydrated particles. Altogether, the input parameters to control electrospun fiber morphology have been characterized with a mechanistic rationale based in particle interaction and elastic rheological behavior. Further, the versatility of this approach needs to be explored, so that the developed method can be useful in a wide range of tissue engineering applications. A variety of decellularization techniques and dECM origins are utilized in dECM scaffolds to meet tissue-specific or disease-specific application requirements. Thus, three common SIS decellularization techniques and three dECM origins were used to prepare dECM electrospinning suspensions and their spinnability was assessed.

### 3.4 Effect of Decellularization Method on Fiber Formation

Decellularized SIS is one of the most widely used dECM materials in clinical practice and current research because of its high growth factor content and proven track record for tissue regeneration.^1, 4^ With its wide use, numerous decellularization techniques have been established to meet different needs due to the potential impact on dECM composition and bioactivity.^52^ The Abraham,^53^ Badylak,^54, 55^ and Luo^6, 56^ decellularization techniques were therefore explored to characterize their impact on SIS suspension properties and spinnability to establish a robust and versatile protocol **(Figure 5)**. SIS was prepared with the Abraham,^53^ Badylak,^54, 55^ and Luo^6, 56^ decellularization techniques. The Abraham technique is a strong chemical treatment involving sodium hydroxide and hydrochloric acid. The Badylak technique is a weak chemical treatment with peracetic acid. In contrast to the Abraham and Badylak chemical only treatment methods, the Luo technique combines an enzymatic digestion treatment with Trypsin-EDTA and SDS and a medium chemical treatment with chloroform, methanol, and peracetic acid. Stronger treatments are more effective at removing cell components, but they also remove some ECM components in the process.^52^ Thus, the proper decellularization method must be chosen based on the application to balance immunogenicity and the preservation of the native ECM for the retention of bioactivity. Due to the impacts of the decellularization on the ECM components, each electrospinning suspension must be tailored for spinnability. Each suspension was prepared with a dried particle size of 500 – 2000 µm. To ensure the suspensions had a viscosity which prevents beading, the Abraham suspension was prepared with a concentration of 40 mg/mL, the Badylak suspension was prepared with a concentration of 60 mg/mL, and the Luo suspension was prepared with a concentration of 70 mg/mL. In this study, an electrospun mesh was fabricated from all three of the decellularization techniques tested, but the suspension properties and electrospun fiber morphology from the Luo technique differed from the Abraham and Badylak techniques **(Figure 5)**. The Abraham and Badylak techniques show more similar particle morphology with similar size and fibril protrusions **(Figure 5A)**. Though, the Badylak suspension has denser particle cores distributed through the looser particle network. This is likely due to the reduced chemical treatment maintaining stronger protein binding that is more difficult to disrupt with homogenization. The Luo suspensions contain larger conglomerates of suspended fibrils on the order of 100 µm. The Luo dECM is likely more homogenous due to increased digestion and processing, so the particles do not reduce energy by separating. Whereas the Abraham and Badylak suspension particles likely break along areas of weaker protein binding innate to a heterogeneous mixture of ECM components. In terms of rheology, the ascending crossover strain amplitude remained consistent across techniques indicating that the ability to resist flowing and breaking under electrostatic forces is similar **(Figure 5D)**. This is supported by the continuous fiber formation seen in each electrospun mesh **(Figure 5B)**. However, the raw dynamic moduli curves with respect to strain amplitude differ **(Figure 5C)**. Though the loss and storage modulus crossover strain amplitude is the same, the negative slope of the dynamic moduli at low strain amplitudes is unique to the Luo suspension. The Abraham and Badylak suspensions exhibit a plateau in the dynamic moduli at low strain amplitudes. This suggests that the particles in the Luo suspension are continuously deforming as strain amplitude is increased until the particle interactions are overcome and the suspension begins to flow. The particles in the Abraham and Badylak suspensions remain static and hold together more consistently until a threshold is reached to separate the particles and initiate flow of the suspension. It is hypothesized that the enzymatic digestion in the Luo technique disrupts protein interaction via denaturation and reduces the binding or attraction force between the particles in suspension. This allows the particles to deform with greater ease under applied strain. When electrospinning, the Luo suspensions are able to deform gradually and more homogenously to form more uniform fibers. The proteins can rearrange more easily under applied strain without completely breaking apart due to the weaker protein interaction and bigger particle conglomerates. These results further support the use of the ascending crossover strain amplitude as a predictor of spinnability. Although the Luo suspension has complex rheological properties that impact fiber morphology differentially compared to the Abraham and Badylak suspensions, the spinnability and fiber formation remains consistent as with the ascending crossover strain amplitude. Additionally, the uniform fiber morphology electrospun from the Luo suspension further reinforces the idea that increased digestion of dECM is beneficial to homogenous processing and scaffold fabrication. However, it is well established that enzymatic digestion denatures proteins and reduces the bioactivity of dECM.^24^ Therefore, there is an interplay between electrospinning scaffolds with homogenous fiber structures and limiting denaturation that cam impact bioactivity.

**Figure 5:**
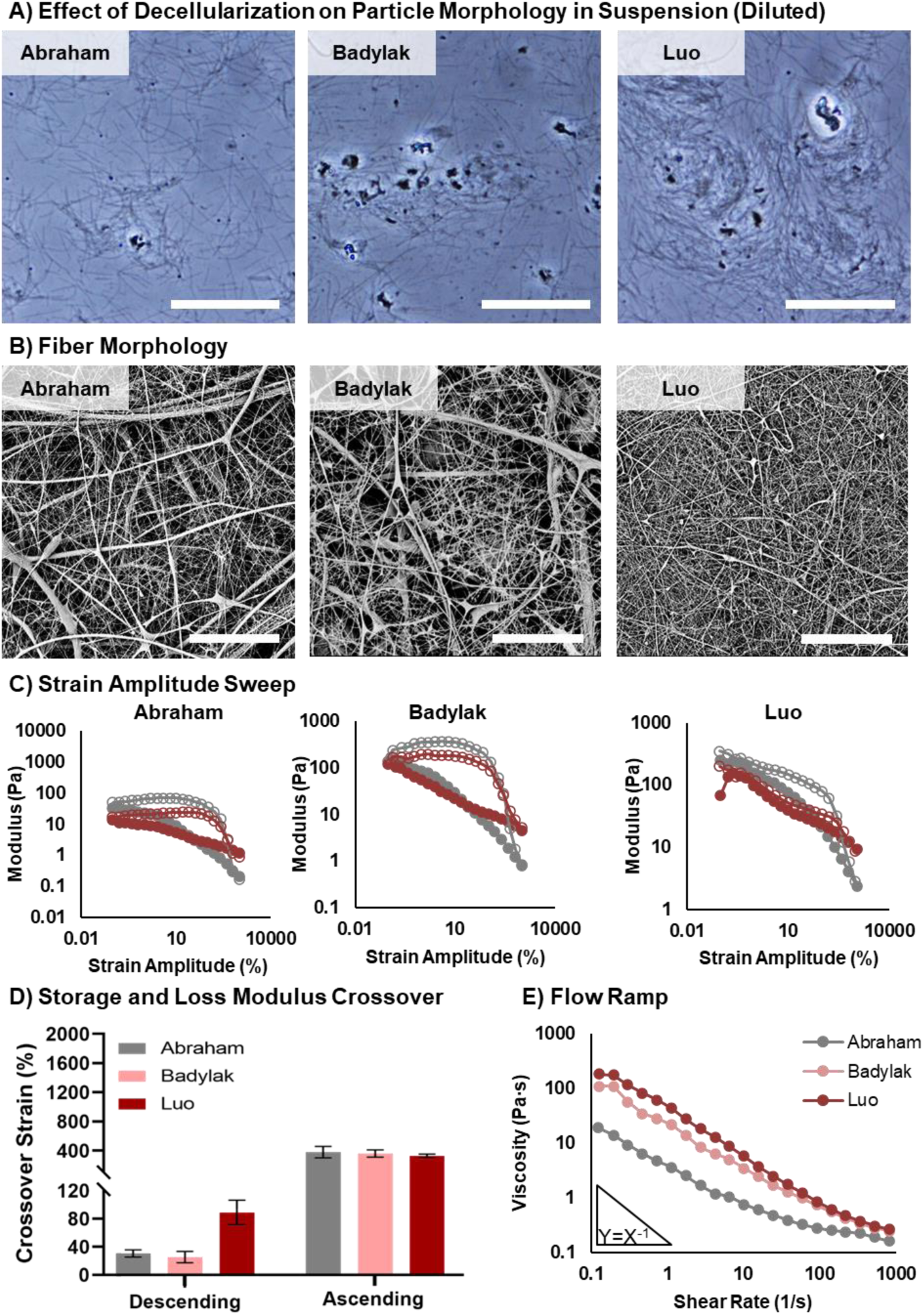
SIS electrospinning utilizing Badylak, Abraham, and Luo decellularization techniques. A) Brightfield image of dECM particles in an electrospinning suspension. Scale bar = 60 µm. B) SEM images of the electrospun dECM fibers. Scale bar = 20 µm. C) Strain amplitude sweep of dECM suspensions in which solid points were collected with a descending strain sweep and hollow points were collected with an ascending strain sweep. D) The crossover strain of the storage and loss modulus curves collected with a descending strain sweep. E) The viscosity with respect to shear rate of the dECM suspensions.

### 3.5 Effect of Tissue Origin on Fiber Formation

The bioactivity of dECM scaffolds can also be controlled by dECM origin. Each tissue has a specialized set of proteins, proteoglycans, and growth factors that are tailored to the functions of the tissue.^70^ For example, SIS and dermal tissues contain a high concentration of growth factors, such as VEGF, TGFβ, and bFGF, to support a constantly regenerating organ. Alternatively, cardiac tissue contains an abundance of fibronectin to maintain homeostasis and recruit myofibroblasts for repair.^4, 70–72^ Tissue engineering scaffolds to treat various tissues and diseases may require a broad range of dECM origin tissues. Thus, the established suspension electrospinning must be functional with many types of dECM. In this study, dECM suspensions were prepared from SIS, skin, and heart tissues **(Figure 6)**. Each suspension was prepared with a dried particle size of 500 – 2000 µm. For adequate decellularization efficiency, the SIS and skin was decellularized with the Abraham technique, and the heart tissue was decellularized with the Badylak technique. To ensure the suspensions had a viscosity which prevents beading, the SIS suspension was prepared with a concentration of 40 mg/mL, the skin suspension was prepared with a concentration of 70 mg/mL, and the Luo suspension was prepared with a concentration of 55 mg/mL. The viscosity flow ramps of the suspensions composed with each dECM origin were similar. The ascending crossover strain amplitude of the SIS and skin suspensions is similar and ∼ 400% while the heart suspension has an ascending crossover strain amplitude significantly lower near 150% **(Figure 6C)**. This indicates that there is reduced particle interaction in the heart-based suspension. Since these suspensions were all prepared with dual homogenization, it is expected that the ECM makeup of heart tissue has reduced binding affinity that reduces the particle interaction forces within the suspension. The load that heart tissue can withstand without fracture is less than SIS and skin, so the ECM intermolecular forces are likely reduced.^73–76^ Once electrospun, a mesh composed of continuous fibers was fabricated with each dECM origin indicating that a crossover strain amplitude of 150% is sufficient to maintain elasticity and withstand the electrostatic forces involved in electrospinning. This further refines the acceptable range of suspension properties for spinnability, where < 50% crossover strain amplitude forms primarily beads and > 100% forms fibers when electrospun. Although these suspensions were all spinnable, their fiber morphology differed. The mesh composed of skin-derived dECM contains localized areas of fusion, and the mesh composed of heart-derived dECM contains flat ribbon-like fibers that are larger than the fibers in the SIS and skin-based meshes. Skin tissue has localized areas of hydrophobic and hydrophilic domains to provide a sufficient external barrier and prevent infection. The varying compositions may phase separate during homogenization and mixing and cause localized areas of altered fiber morphology. For the heart mesh, ribbon formation has been previously explained as reduced drying of the solvent leading to fiber collapse and/or the reduced stretching or elasticity of the solution during drawing.^49^ Since each suspension contains the same solvent and spinning conditions, it is unlikely that the evaporation rate of HFIP during electrospinning is different between the SIS, skin, and heart suspensions. However, the reduced crossover strain amplitude of the heart suspension indicates a reduction of elasticity of the suspension at higher strains. Thus, the suspension is drawn less under electrostatic forces and stays in bigger fibers that dry quickly on the outside and collapse before the inner fiber can completely dry and retain its shape.

**Figure 6:**
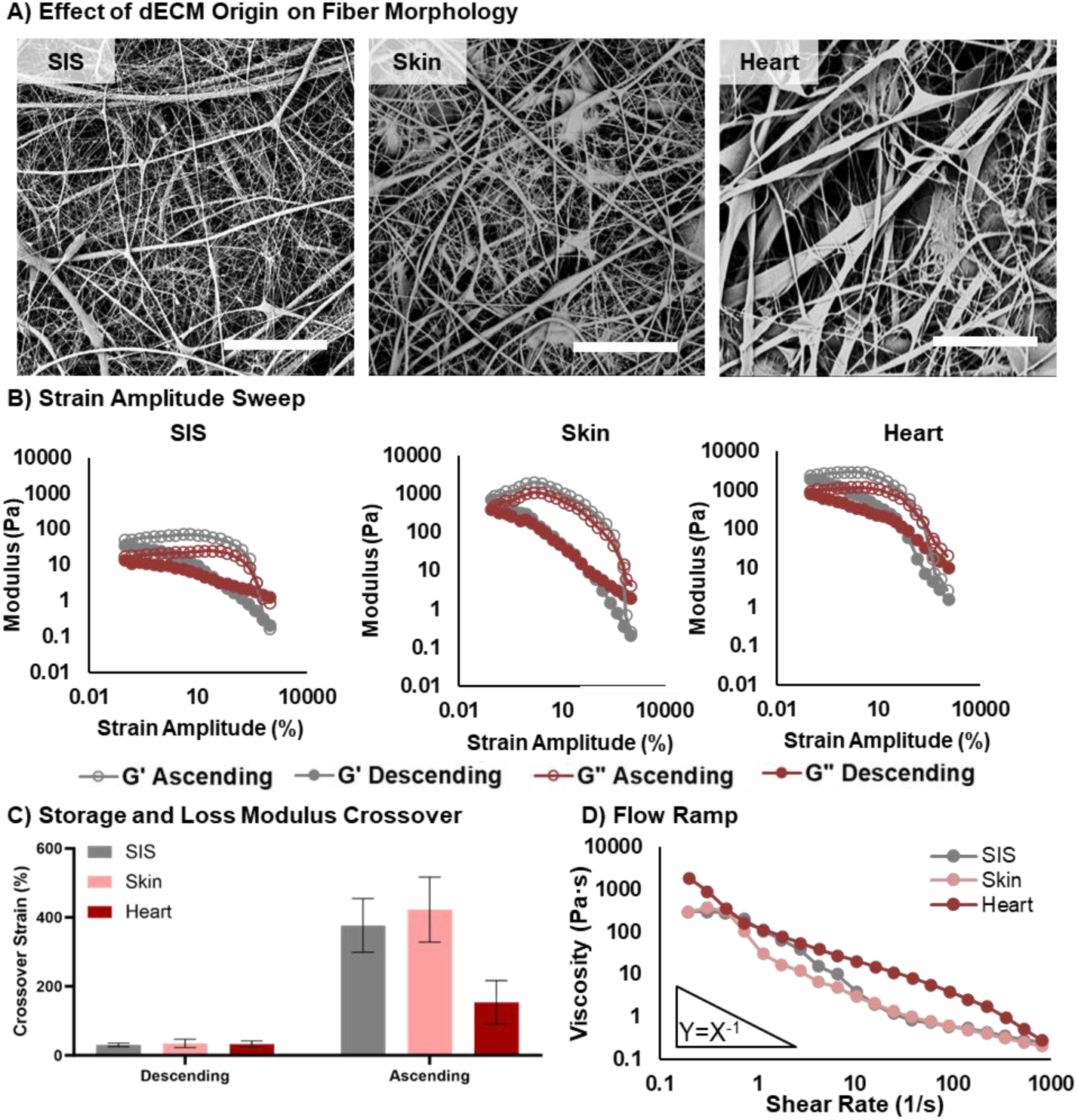
Suspension electrospinning of SIS, skin, and heart-derived dECM. A) SEM images of the electrospun dECM fibers. Scale bar = 20 µm. B) Strain amplitude sweep of dECM suspensions in which solid points were collected with a descending strain sweep and hollow points were collected with an ascending strain sweep. C) The crossover strain and modulus of the storage and loss modulus curves collected with a descending strain sweep. D) The viscosity with respect to shear rate of the dECM suspensions.

Herein, the versatility and robustness of the dECM suspension electrospinning technique is showcased and a mechanistic rationale based in suspension rheology and dECM composition is used to explain the differences in electrospun fiber morphology. Enzymatic digestion can be used to create more homogenous fibrous scaffolds, but this could reduce the bioactivity of the scaffold via denaturation. Additionally, dECM origin will impact the suspension rheological properties and fiber morphology of the electrospun mesh based on different compositions, but the suspensions that maintain a crossover strain amplitude above 100% have been successfully electrospun. Since the dECM suspensions retain protein microstructure above the tertiary organization level, as indicated by fibrillar components on the same scale as collagen bundles, it is hypothesized that the electrospun dECM meshes retain the bioactivity of native dECM. In order to test this hypothesis, cell proliferation, angiogenic bioactivity, and immunomodulatory bioactivity before and after electrospinning the decellularized SIS was assessed **(Figure 7)**.

**Figure 7:**
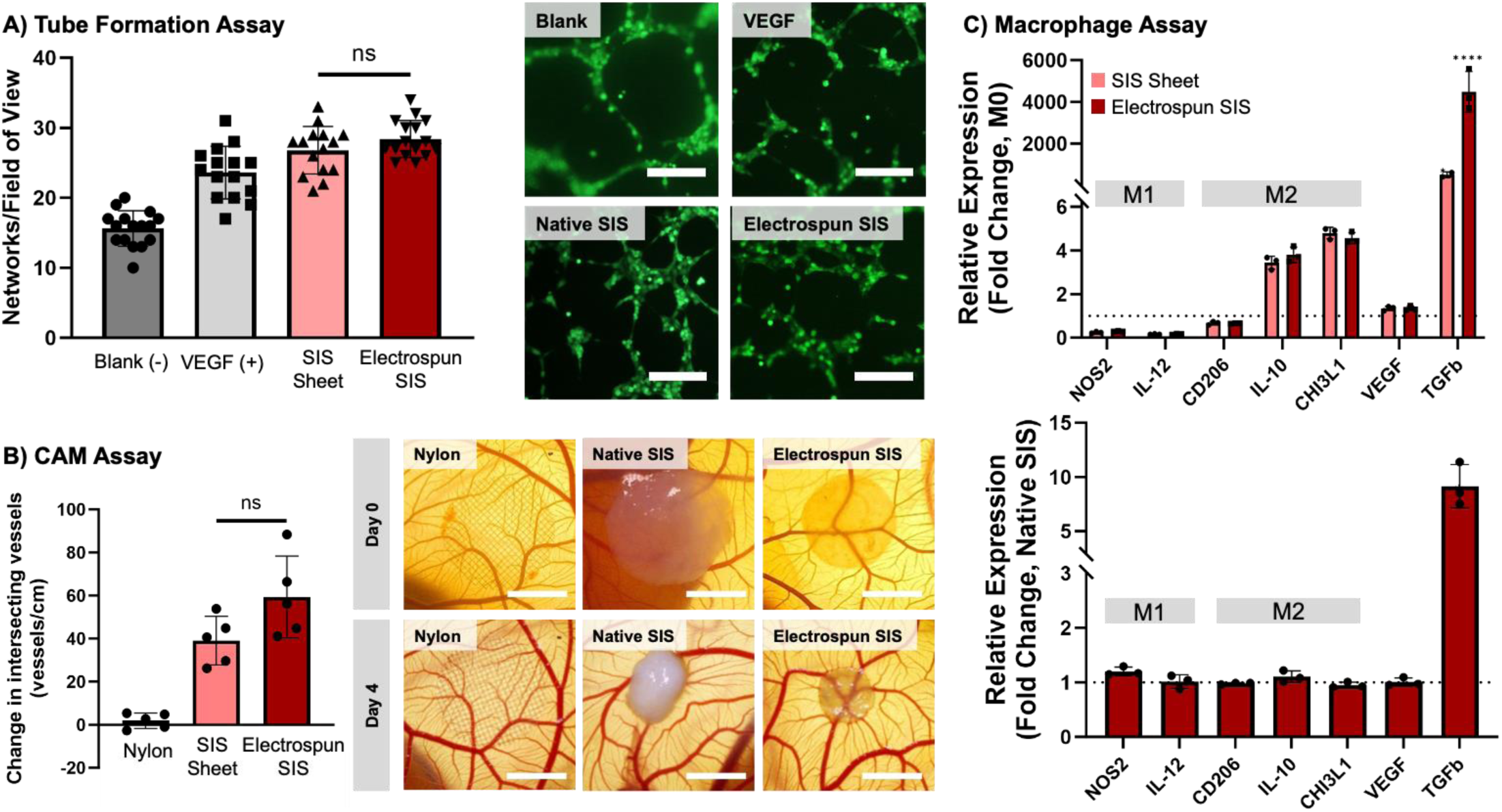
Evaluation of the retention of bioactivity after the electrospinning process. A) *In vitro* HUVEC tube formation in SIS-conditioned media. Scale bar = 500 µm. B) *In ovo* vascularization in a CAM assay after 4 days of sample placement. Vessel density is calculated as the change in the number of vessels intersecting with the sample from Day 0 to Day 4, normalized to sample perimeter. Scale bar = 3 mm. C) Macrophage gene expression after 72 h of SIS contact with respect to the expression of the M0 control (top) and native SIS (bottom).

### 3.6 Bioactivity Retention of Electrospun dECM

Cell proliferation was assessed as an initial measure of the biocompatibility of the electrospun SIS. Human dermal fibroblasts were cultured on each substrate and the metabolic activity of the cells were measured using an MTS assay after 1, 3, and 7 days. The metabolic activity of the cells cultured on each substrate increased over time with no significant difference between the SIS sheet and the electrospun SIS mesh **(Figure S1)**. This indicated that the electrospinning process did not have a negative impact on cell attachment and proliferation to the SIS substrate.

The angiogenic capacity of SIS has been widely studied and is one of the primary considerations when choosing SIS as a regenerative scaffold material.^5, 6, 77–79^ Vascularization and nutrient support is key to supporting tissue regeneration. Thus, it is important that the angiogenic capacity of SIS was retained after electrospinning. An in vitro tube formation study and an *in ovo* chorioallantoic membrane (CAM) assay were both conducted to robustly assess the impact of electrospinning on SIS angiogenic bioactivity. For the tube formation study, HUVECs were cultured on Reduced Growth Factor Matrigel® in SIS-conditioned media for 4 hours **(Figure 7A)**. The number of capillary-like networks formed in response to the conditioned media was compared between a blank control, a positive VEGF control (20 ng/mL), a SIS sheet, and an electrospun SIS mesh. All media groups supported some level of tube formation due to the angiogenic cues in Matrigel. However, the increase in networks that formed in response to the positive VEGF control compared to the negative blank control indicated that the network-forming capacity of the HUVECs was not saturated. Further, both SIS substrates supported elevated network formation compared to the blank control and are similar to the VEGF control. Also, the electrospun SIS-conditioned media supported similar network formation to the SIS sheet-conditioned media. This indicates that the electrospinning process did not negatively impact the angiogenic capacity of the SIS with regards to release products. Since the tube formation assay is limited to the assessment of release products and is artificially inflated by the Matrigel, a CAM assay was conducted for a more robust analysis of the SIS angiogenic capacity. After 10 days of quail embryo culture, a nylon mesh (negative control),^80^ SIS sheet, and electrospun SIS were placed on the CAM for an additional 4 days **(Figure 7B)**. The change in vasculature was assessed over time with daily imaging. After 4 days of treatment, the primary change in vasculature expected to be seen is small capillary-like vessels, so the vessel density was quantified as the number of vessels intersecting the sample perimeter instead of a vessel area calculation that is heavily skewed by large vessels and sample placement. The Nylon mesh did not significantly impact the vessel density after 4 days on the CAM, indicating that angiogenic cues are needed to increase the vasculature within this timeframe, and merely agitation or sample placement on the membrane does not impact angiogenesis. Both of the SIS substrates supported elevated vessel formation compared to the Nylon mesh, and the electrospun SIS mesh supported similar levels of vessel formation compared to the SIS sheet. Both of the SIS substrates also reduce in size after 4 days on the CAM meaning cells likely migrated into the samples and contracted the ECM. Together, the tube formation assay and CAM assay support that the angiogenic capacity of the SIS is retained and not negatively impacted after being electrospun.

Lastly, the immunomodulatory bioactivity of the electrospun SIS was evaluated via macrophage polarization. The macrophage response to a material can control the wholistic biological response including dictating inflammatory versus regenerative behavior.^81, 82^ For this reason, THP1-derived macrophages were cultured on the decellularized SIS sheet, electrospun SIS mesh, and a negative PDMS-coated well control. The gene expression of the macrophages after 72 hours of culture of NOS2, IL12, CD206, IL10, CHI3L1, VEGF, and TGFβ was measured via qPCR, **Figure 7C**. The SIS substrates induced a reduction in NOS2 and IL12 expression and increased expression of IL10, CHI3L1, and TGFβ compared to the M0 negative control. This indicated the macrophages cultured on the SIS substrates have a preference toward tolerogenic polarization. The expression of macrophages cultured on the electrospun SIS mesh was largely similar to the SIS sheet, with statistically similar (p < 0.05) expression of NOS2, IL12, CD206, IL10, CHI3L1, and VEGF. The only gene impacted differentially by the electrospun scaffold is TGFβ, where there was a 10-fold increase in TGFβ expression between the electrospun SIS mesh and the SIS sheet. It is hypothesized that the physical structure and morphology of the electrospun SIS mesh is impacting the expression profile because the 72 hour exposure to SIS is not expected to be long enough for the impacts of differential gene expression. TGFβ plays a role in tissue remodeling by supporting tissue regeneration without scar formation and increasing expression of IL10 and CD206 in macrophages.^83, 84^ Electrospinning did not largely impact the immunoregulatory bioactivity of SIS via macrophage gene expression with the exception of TGFβ and supports a tolerogenic and anti-inflammatory immune response. Altogether, the angiogenic and immunomodulatory bioactivity of SIS was not significantly impacted by suspension electrospinning. This is largely attributed to the lack of enzymatic digestion and additives required in processing and the retention of protein microstructure above the tertiary organization level.

## 4. Conclusion

In this study, we conducted a rigorous evaluation of dECM suspension properties necessary to support fiber formation during electrospinning. Homogenization was used as a mechanism to increase particle interaction and elastic behavior of the dECM suspensions. The crossover strain amplitude of the loss and storage modulus curves above 100% has been identified as a functional rheological predictor of a spinnable dECM suspension. The sensitivity of fiber morphology to suspension concentration was shown to be similar to polymer solution concentration, and interestingly, the dried particle size range tested broke down to a consistent hydrated state and electrospun into a similar fiber morphology. The versatility of this dECM suspension electrospinning approach was showcased by utilizing three different decellularization techniques and three different dECM tissue origins with varying compositions. These groups further supported the need for a crossover strain amplitude above 100% and highlighted the impact of enzymatic digestion on dECM processing. With the variables tested, the impact of each of the steps to prepare a dECM suspension on electrospinning were evaluated, so the protocol can be tuned for a specific application with full knowledge of the influence on the resulting electrospun scaffold. The motivation for suspension electrospinning is to increase the retention of bioactivity after scaffold processing that is commonly lacking with the addition of a carrier polymer or enzymatic digestion. The regenerative capacity of an electrospun SIS scaffold was shown to be similar to a decellularized SIS sheet with regards to cell proliferation, angiogenesis, and immunomodulation. This study elucidated the critical parameters that guide dECM suspension electrospinning to provide researchers with a framework to use this new approach to create dECM scaffolds that retain its native regenerative capacity.

