## Supplemental Figure 1 for "Suspension Electrospinning of Decellularized Extracellular Matrix"

SUPPLEMENTAL INFORMATION:

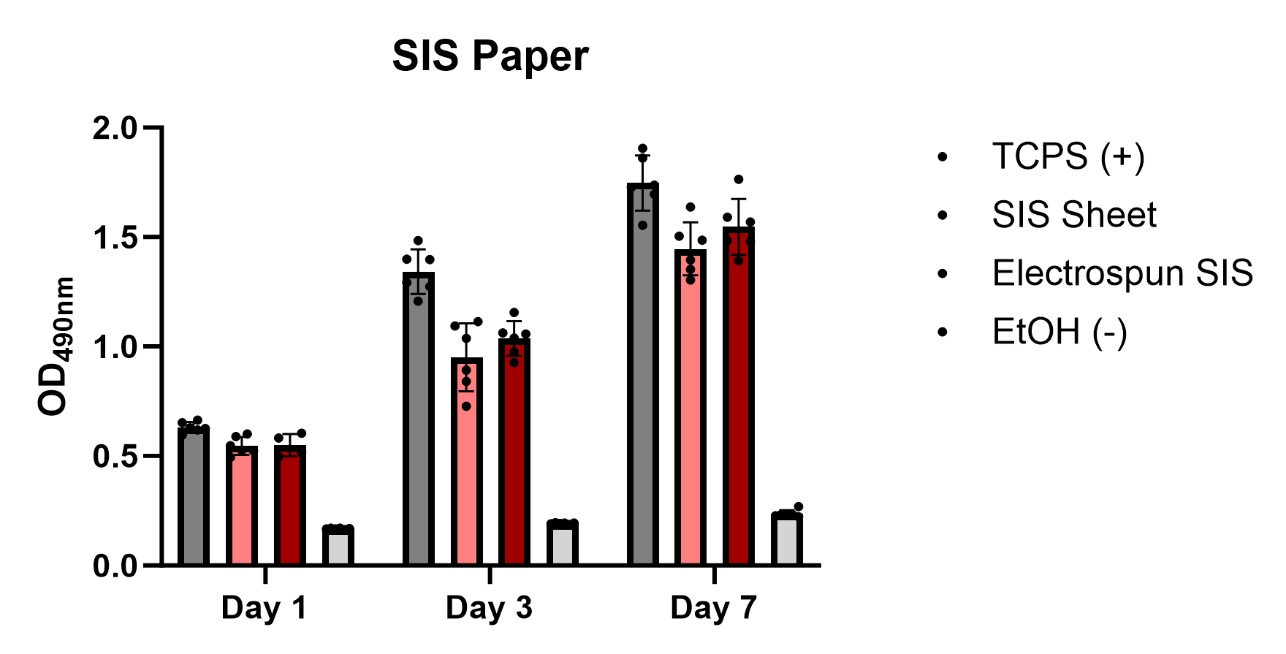

SIS Sheet

TCPS (+)

Electrospun SIS

EtOH (-)

***Figure S1:*** Evaluation of the retention of bioactivity after the electrospinning process. Cell proliferation of human dermal fibroblasts on SIS substrates.
